# CCPE: Cell Cycle Pseudotime Estimation for Single Cell RNA-seq Data

**DOI:** 10.1101/2021.06.13.448263

**Authors:** Jiajia Liu, Mengyuan Yang, Weiling Zhao, Xiaobo Zhou

## Abstract

The rapid development of single-cell RNA-sequencing (scRNA-seq) technologies makes it possible to characterize cellular heterogeneity by detecting and quantifying transcriptional changes at the single-cell level. Pseudotime analysis enables to characterize the continuous progression of various biological processes, such as cell cycle. Cell cycle plays an important regulatory role in cell fate decisions and differentiation and is also often regarded as a confounder in scRNA-seq data analysis when analyzing the role of other factors on transcriptional regulation. Therefore, accurate prediction of cell cycle pseudotime and identify cell stages are important steps for characterizing the development-related biological processes, identifying important regulatory molecules and promoting the analysis of transcriptional heterogeneity. Here, we develop CCPE, a novel cell cycle pseudotime estimation method to characterize cell cycle timing and determine cell cycle phases from single-cell RNA-seq data. CCPE uses a discriminative helix to characterize the circular process and estimates pseudotime in the cell cycle. We evaluated the model performance based on a variety of simulated and real scRNA-seq datasets. Our results indicate that CCPE is an effective method for cell cycle estimation and competitive in various downstream analyses compared with other existing methods. CCPE successfully identified cell cycle marker genes and is robust to dropout events in scRNA-seq data. CCPE also has excellent performance on small datasets with fewer genes or cells. Accurate prediction of the cell cycle in CCPE effectively contributes to cell cycle effect removal across cell types or conditions.

## Introduction

The rapid development of single-cell RNA-sequencing (scRNA-seq) technologies makes it possible to characterize cellular heterogeneity in gene expression at single-cell resolution [1–4]. Cell cycle is a fundamental component in the biological processes and the main driver of transcriptional heterogeneity [5, 6]. During development/embryogenesis, embryo stem cells/progenitor cells undergo self-renewal and lineage-specific differentiation programs to generate specific cell types. In adulthood, stem cells continue to differentiate and create fully differentiated progeny cells during tissue repair and normal cell renewal. Cell cycle plays an important regulatory role in stem cell fate decisions [7] and differentiation [8]. As the main rate-limiting step of differentiation [8], cell cycle control is essential in ensuring generating cell diversity and maintaining the homeostasis of adult tissues. Cancer cells are derived from cancer stem cells/progenitor cells and can also differentiate from cancer cells to re-enter the cell cycle and become cancer progenitor cells [9, 10]. Loss of cell cycle control can lead to uncontrolled tumor cell proliferation and growth [11]. In addition to the significance in studies of tumorigenesis and development [12–14], the cell cycle is often regarded as a confounder in scRNA-seq data analysis when analyzing the role of other factors on transcriptional regulation. Removing confounder effect will improve the resolution of other biological processes [15]. Therefore, accurately identifying the cell cycle of individual cells is needed to fully understand a number of different biological problems and data analysis issues.

Many experimental methods, such as utilizing chemical induction [16], counterflow centrifugation elutriation [17] and DNA content [18] have been used to detect the cell cycle phases of individual cells [19]. However, these methods are time-consuming and laborious, and not a quantitative measurement of cell cycle phase duration. Nowadays, more and more computational methods are developed to address the disadvantages in experimental cell cycle estimation, including knowledge-based and unsupervised approaches. Knowledge-based methods, such as cyclone [6] and CellCycleScoring function in Seurat [20], use annotated cell cycle genes to predict the classification of each cell in G1, S or G2/M phase. Peco is another knowledge-based approach that uses the data generated from FUCCI fluorescence images and scRNA-seq to train the “naive Bayes” predictor for predicting the progression position of each cell through the cell cycle process, which we called cell cycle pseudotime [21]. Peco is specially designed for human induced pluripotent stem cells. reCAT requires cell cycle marker genes to calculate the Bayes-scores of each cell, and uses consensus traveling salesman problem (TSP) and hidden Markov model (HMM) to recover cell cycle pseudo time-series and stages [22]. Such knowledge-based methods have a common drawback, that is, their applications are limited to datasets with pre-annotated cell cycle genes and experimental cell cycle labels. To address this problem, several unsupervised methods have been proposed to predict cell cycle pseudotime, such as Cyclum [23] and CYCLOPS [24]. Cyclum employs an autoencoder model that takes both non-linear and linear components in the hidden layer into account. The non-linear projection of gene expression profiles is trained to infer the pseudotime of cells in the circular process [23]. CYCLOPS uses an autoencoder model with linear projection to project data onto a closed elliptical curve in low-dimensional space [24]. However, CYCLOPS employs square root and division in the autoencoder model, which makes optimization more complicated.

In this study, we proposed a novel unsupervised method named CCPE to estimate cell cycle pseudotime of single cells from single-cell RNA-seq data. CCPE learns a discriminative helix to characterize the circular process and estimates pseudotime in the cell cycle. We assessed the performance of CCPE in estimating cell cycle pseudotime and stage assignment by applying CCPE to several downstream analyses using both simulated and real scRNA-seq datasets. We also assessed the performance of CCPE in handling dropout events, analyzing smaller datasets with fewer genes or cells and removing cell cycle effect from scRNA-seq data.

## Results

### Overview of CCPE approach

Single-cell RNA sequencing (scRNA-seq) data is a cell-specific gene expression matrix with high dimensionality and sparsity. Traditional clustering methods have low efficiency for computing high-dimensional and sparse matrices. Therefore, it is necessary to introduce dimension reduction in the model. We develop CCPE, a novel cell cycle pseudotime estimation method to characterize cell cycle timing from single-cell RNA-seq data. CCPE maps high-dimensional scRNA-seq data onto a helix in three-dimensional space, where 2D space is used to capture the cycle information in scRNA-seq data, and one dimension to predict the chronological orders of cells along the cycle, which we called cell cycle pseudotime. ScRNA-seq data is repeatedly transformed from high dimensional to low dimensional and then mapped back to high dimensional. At the same time, CCPE iteratively optimizes the discriminative dimensionality reduction via learning a helix until convergence (**Figure 1**). CCPE is applied to several downstream analyses and applications to demonstrate its ability to accurately estimate the cell cycle pseudotime and stages.

**Figure 1.**
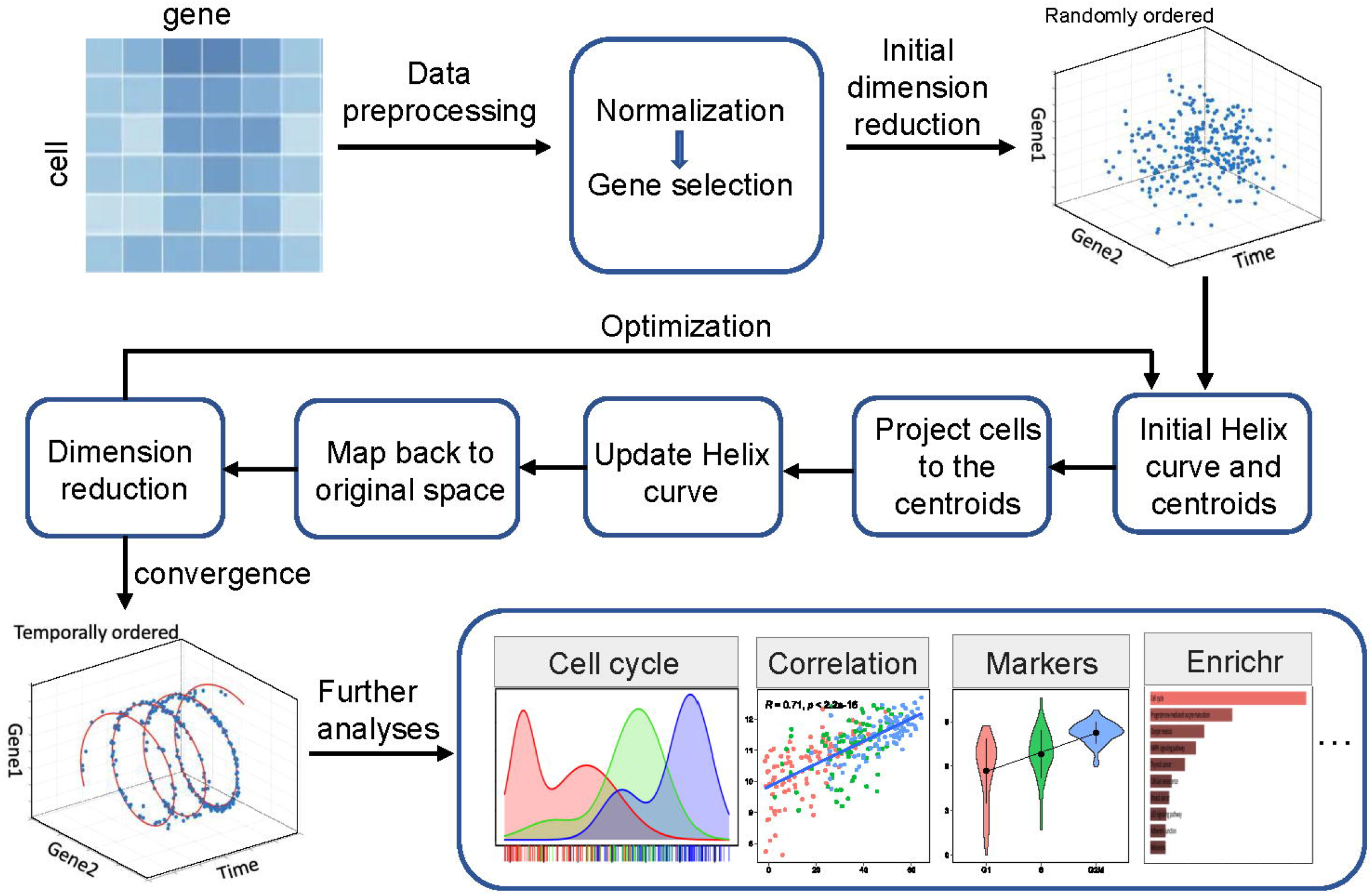
Overview of CCPE approach. After normalization and pre-processing of the data, CCPE learns the discriminative helix by iteratively optimization between the original and reduced dimension until convergence. After optimization, a 3D-helix with two-gene dimensions is used to represent circular information of cell cycle phases and one dimension to represent pseudotime of cells along the cell cycle. Several downstream analyses and applications of CCPE are used to assess its performance.

### Estimation of cell cycle pseudotime

As we mentioned in the introduction, few computational tools have been developed so far to be used for the estimation of cell cycle pseudotime for single cells, including Cyclum, CYCLOPS and reCAT [22–24]. To test the performance of CCPE in predicting the cell cycle pseudotime, we compare the performance of CCPE with Cyclum and CYCLOPS based on scRNA-seq data of mouse embryonic stem cells (mESCs) sequenced by Quartz-Seq technology [18]. **Figure 2a** illustrates the distribution of cell cycle pseudotimes estimated by each method. Both CCPE and Cyclum can maintain the correct cell cycle order from G1 to S, and then to G2/M. Both of CYCLOPS and reCAT can distinguish G1 and S phases well but do not characterize G2/M phase correctly. Compared with Cyclum, CCPE shows a better performance in separating S and G2/M phases. Overall, CCPE has an excellent performance in accurately predicting cell cycle pseudotime of mESCs scRNA-seq dataset.

**Figure 2.**
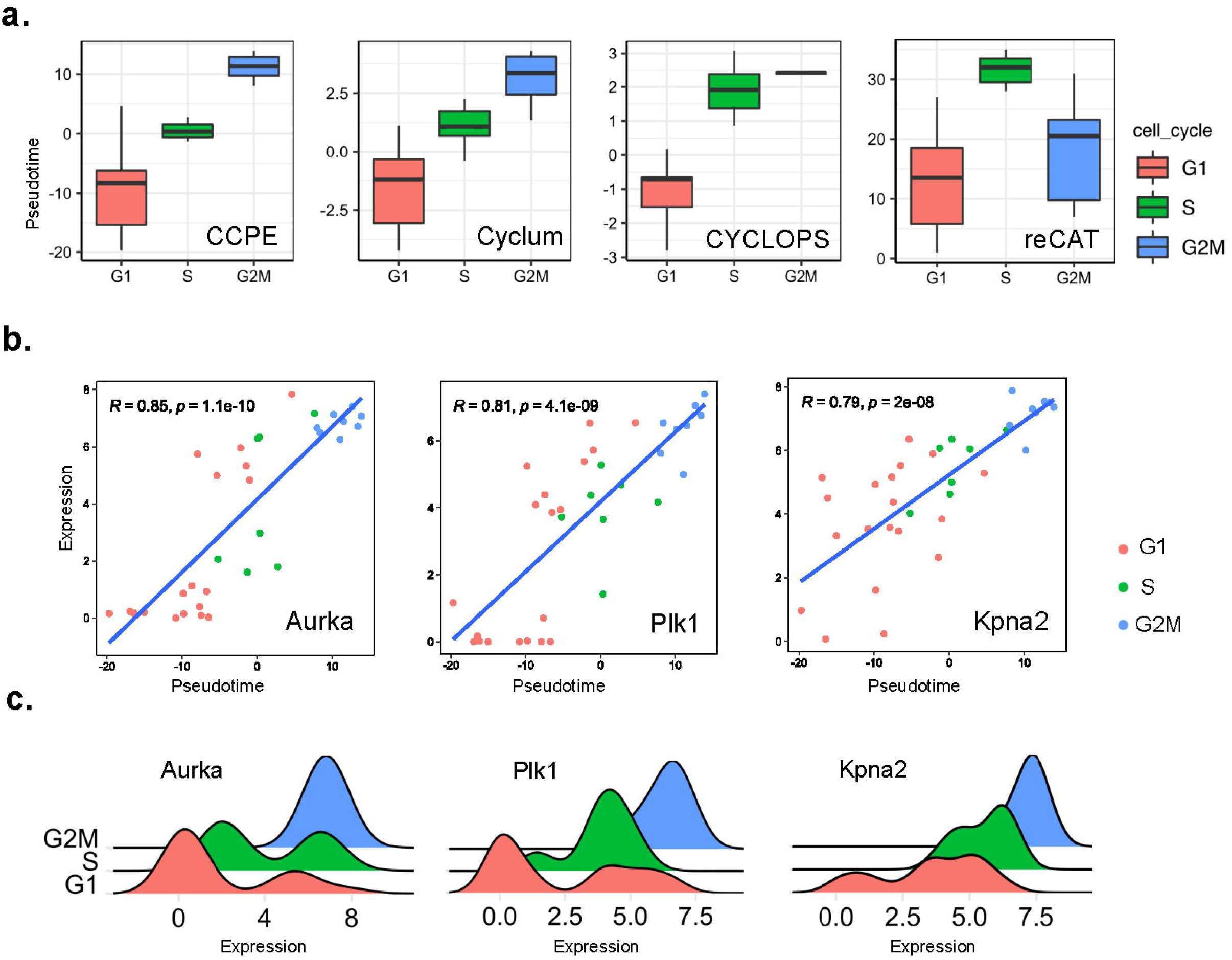
Cell cycle pseudotime analysis of mESCs Quartz-Seq data. **(a)** Boxplots demonstrated the distribution of cell cycle pseudotimes inferred by CCPE, Cylum, CYCLOPS and reCAT, respectively. Each box was colored by the corresponding cell cycle phases and outliers were ignored in the figure. **(b)** The expressions of three cell cycle marker genes are highly correlated with the cell cycle pseudotime estimated by CCPE. The correlation coefficients and p-value are shown on the top left of each figure. **(c)** shows density plots of the expressions of three G2/M-phase marker genes.

We calculated the Pearson correlation of the gene expression and cell cycle pseudotime inferred by CCPE. Aurora kinase A (Aurka), polo-like kinase 1 (Plk1) and karyopherin alpha 2 (Kpna2) have the highest correlation with cell cycle pseudotime. The correlation coefficients of Auraka, Plk1 and Kpna2 genes are 0.85, 0.81, and 0.79, respectively (**Figure 2b**). Aurka is known as a key cell-cycle regulator, whose levels of mRNA and protein are low in G1 and S and increase sharply during G2/M phase [25]. Plk1 has a crucial role in the regulation of mitotic checkpoints and is active in the late G2 phase [26]. Knocking-down Kpna2 has been shown to inhibit cell proliferation by inducing cell cycle arrest in G2/M phase [27]. We found that the most highly correlated genes with cell cycle pseudotime are G2/M-phase marker genes, which are all highly expressed in G2/M phase (**Figure 2c**).

### Assignment of cell cycle stages

We compared the competence in assigning cells into the correct cell cycle stages of CCPE with others models. To do so, we took advantage of a Gaussian mixture model with three components to transform the continuous pseudotime generated by CCPE, Cyclum and CYCLOPS into discrete G1, S and G2/M stages. In addition to Cyclum and CYCLOPS, we also compared CCPE with cyclone, Seurat and reCAT using both mESCs Quartz-Seq and E-MTAB-2805 mESCs datasets. Ten clustering metrics was used to evaluate the models’ performance. Due to the randomness in the model of Cyclum, CYCLOPS, cyclone and reCAT. Each method was evaluated ten times on each dataset and the average values of ten clustering metrics is recorded. The performance of CCPE on mESCs Quartz-Seq is outstanding, having the highest accuracy among all the methods (**Figure 3a**). CCPE also ranks first in all the considered metrics on E-MTAB-2805 mESCs dataset (**Figure 3b**). Knowledge-besed method, cyclone has good performance second to CCPE, but cannot calculate NMI. The overall performance of Cyclum is better than Seurat, CYCLOPS and reCAT, but not as good as CCPE. Even the best values of cyclone and CCPE in each matric are smaller than those of CCPE. Our results demonstrate the excellent performance of CCPE in predicting the cell cycle stages.

**Figure 3.**
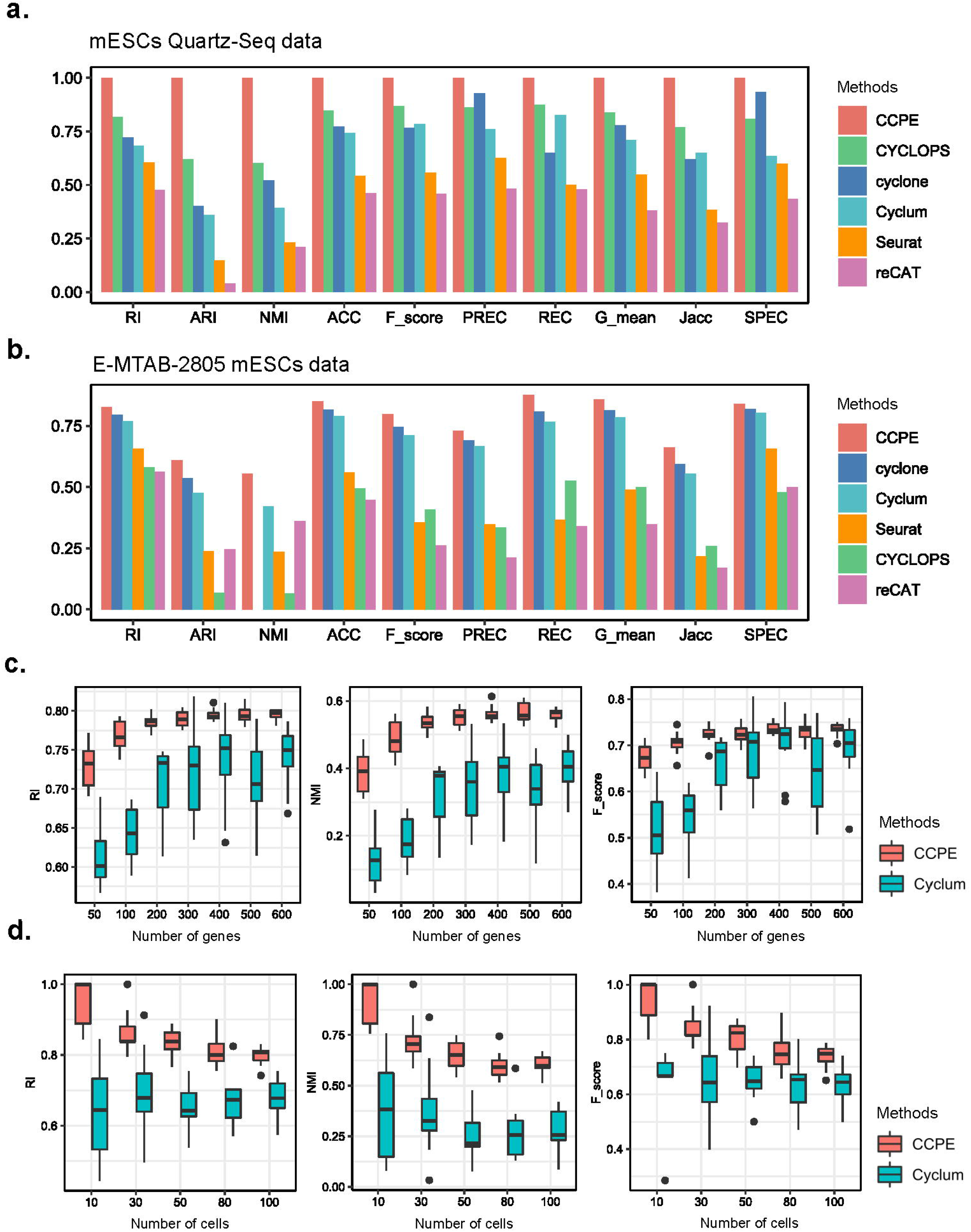
Cell-cycle stage inference from real datasets. **(a)** shows ten clustering measurements to evaluate the cell cycle classification accuracy of CCPE, cyclone, Seurat, reCAT, Cyclum and CYCLOPS for mESCs Quartz-Seq data. **(b)** shows ten clustering measurements to evaluate the cell cycle classification accuracy of CCPE, cyclone, Seurat, reCAT, Cyclum and CYCLOPS for E-MTAB-2805 mESCs data. **(c)** Boxplots of RI, NMI and F_score values indicate the performance of CCPE and Cyclum on the subsampled datasets with smaller number of genes. **(d)** Boxplots of RI, NMI and F_score show the cell cycle classification accuracy of CCPE and Cyclum on the subsampled datasets with different numbers of cells.

### Robustness of CCPE in analyzing small size of scRNA-seq data

To evaluate the performance of CCPE on the data with different numbers of genes and cells, especially the data with a small number of genes and cells, we randomly subsampled the scRNA-seq data from human embryonic stem cell dataset, which consists of 247 cells and 19084 genes. We selected seven subdatasets with different number of genes, ranging from 50 to 600 genes, and five subdatasets with different cell numbers, ranging from 10 to 100 cells. We found that the median of all the clustering metrics of both CCPE and Cyclum gradually increased with the number of genes (**Figure 3c)**. However, when the number of genes increased to more than 300, CCPE consistently outperformed Cyclum in terms of RI, NMI, F_score, ARI, REC and Jacc values. In other words, CCPE can predict cell cycle stages more accurately on a smaller number of genes than Cyclum. CCPE also has better performance on a smaller number of cells compared with Cyclum. The performance of CCPE gradually declines as the number of cells increases and finally stabilizes (**Figure 3d)**. The median value of Cyclum oscillates within a certain range (between 0.65 and 0.68 in RI, between 0.23 and 0.4 in NMI and between 0.63 and 0.66 in F_score), but lower than CCPE. Our analysis indicates that CCPE is more robust and has a higher prediction accuracy for the small datasets with a smaller number of genes or cells.

### Differential gene expression analysis based on inferred cell cycle phases

Differential gene expression analysis of inferred cell cycle phases can identify gene expression variability between different cell cycle phases. We use DESeq2 [28] implemented in R/Bioconductor to detect differentially expressed genes (DEGs) for E-MTAB-2805 mESCs data. DEGs were detected using DESeq2 from CCPE-inferred and Cyclum-inferred cell cycle stages (p.adjusted ≤ 0.05 and |log2FC| ≥ 1). The CCPE-DEGs were obviously involved in the cell cycle pathways and enriched in the biological cell cycle-related processes, including p53 signaling pathway, progesterone-mediated oocyte maturation and circadian rhythm. However, pathways enriched by Cyclum-DGEs have little relationship with the cell cycle (**Figure 4a)**. **Figure 4b** shows the expression of three G2/M phase marker genes Plk1, Bub3, Cdc20 and Fzr1, which are enriched in the cell cycle pathway.

**Figure 4.**
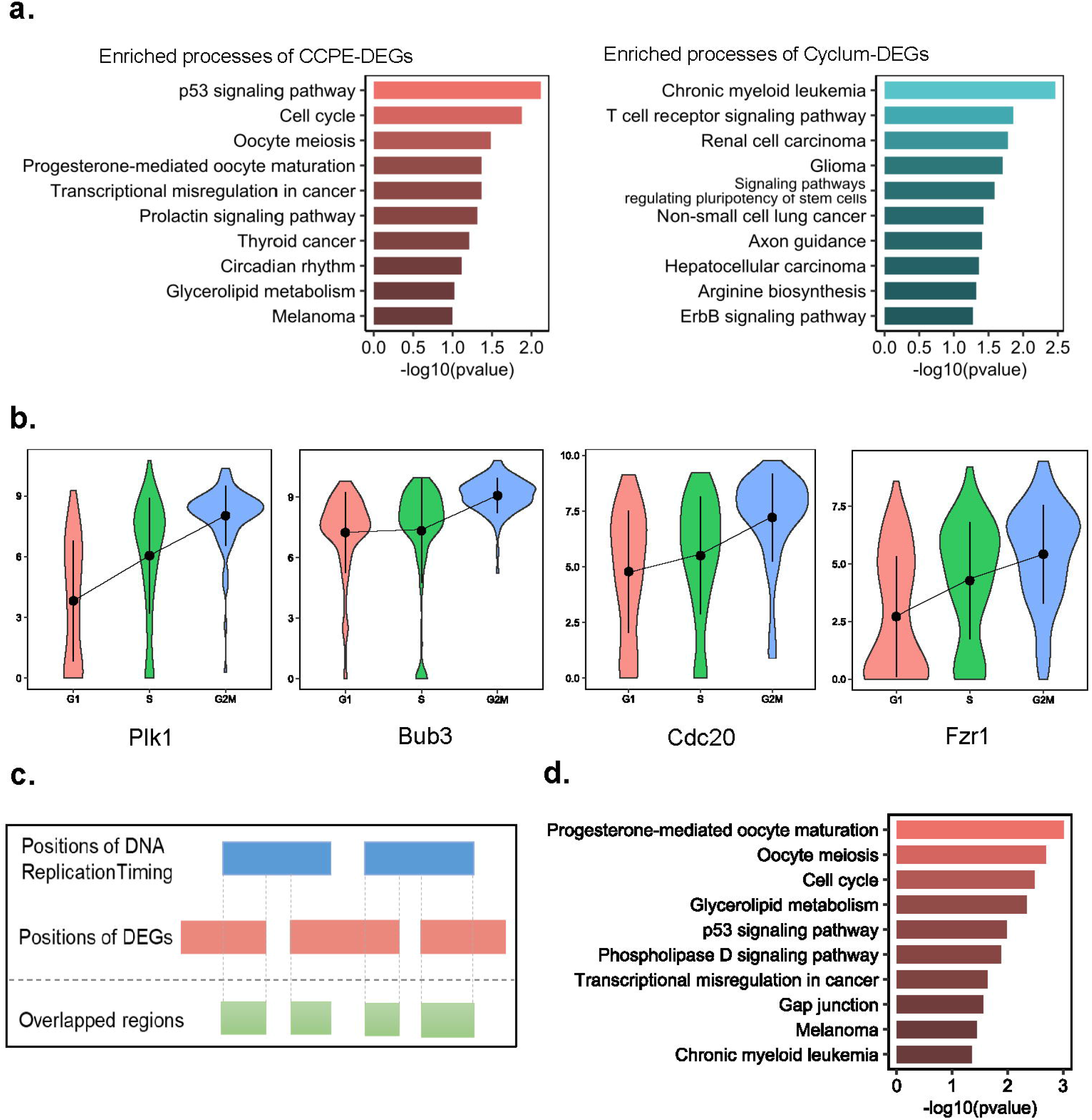
CCPE-inferred cell cycle pseudotime and differentially expressed gene analysis on E-MTAB-2805 mESCs data. **(a)** Top ten enriched biological progresses associated with DEGs identified by CCPE-inferred and Cyclum-inferred cell cycle stages. **(b)** The violin plots show the expression of G2/M phase marker genes Plk1, Bub3, Cdc20 and Fzr1. **(c)** Schematic representation of the intersection between DEGs and DNA replication timing events. **(d)** Top ten enriched biological progresses of the 682s DEGs overlapped with DNA replication timing events.

To further understand the functions of these differential genes in the cell cycle, we intersected the positions of 1577 CCPE-DEGs on the chromosomes with DNA replication timing events of the human genome using *intersect* function in BEDTools [29] (**Figure 4c**). The DNA replication timing program of the cell is highly organized and defined as the temporal sequence of locus replication events during the S phase of the cell cycle [30, 31]. We found that 682 out of 1557 DEGs are overlapped with features in DNA replication timing. Gene ontology (GO) enrichment analysis [32] of the overlapped genes shows the enriched GO terms are mainly associated with the regulation of the cell cycle processes **(Figure 4c)**. Overall, CCPE can not only provide an accurate estimation of cell cycle stages, but also improve the identification of differentially expressed genes and facilitate the search for genes that regulate the DNA replication timing.

### Robustness of CCPE in dealing with dropout events of scRNA-seq data

scRNA-seq data always suffers from many sources of technical noises, leading to excess false zero values, which are termed as dropout events [33]. The tools developed for analyzing scRNA-seq data should take their ability to handle dropout events into account. We used three simulated datasets with different dropout rates (25.6%, 51.1% and 68.8%) generated by *scSimulator* function in CIDR [34] to evaluate the robustness of CCPE in dealing with dropout events. **Figure 5a** shows the UMAP visualization of simulated cells at different CCPE-inferred cell cycle stages. As the dropout rate increases, the performance of CCPE on separating three cell cycle clusters gradually decreases. When the dropout rates are 51.1% and 68.8%, it is difficult for CCPE to distinguish the three cell cycle phases **(Figure 5a)**. As we all know, the higher the dropout rate, the more gene expression are lost. We compared the impact of drop-out rate on the performance of CCPE with Cyclum and CYCLOPS. Through the calculation of clustering evaluation metrics **(Figure 5b)**, we can see that when the dropout rate is less than 51.1%, CCPE performs significantly better than Cyclum and CYCLOPS. When the dropout rate increases to 68.8%, CCPE, Cyclum and CYCLOPS all performed poorly in estimating cell cycle phases. The RI, ARI, NMI, ACC, G_mean and PREC of CCPE are still higher than Cyclum and CYCLOPS. Thus, our analysis results indicate that CCPE is more robust to dropout events than Cyclum and CYCLOPS.

**Figure 5.**
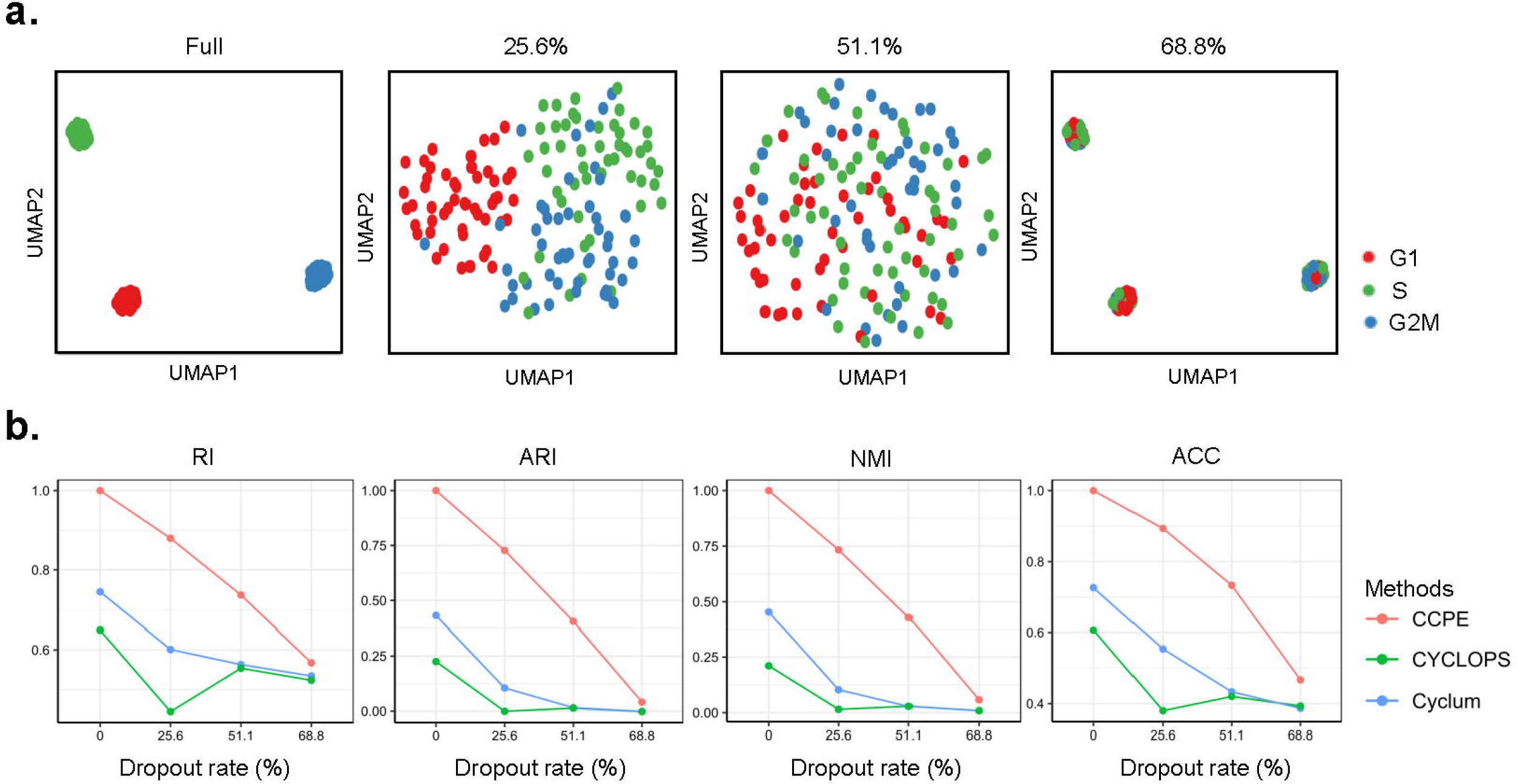
Robustness of CCPE in dealing with dropout events in simulated dataset. **(a)** UMAP plots for data with 0%, 25.6%, 51.1% and 68.8% dropout rate. Full represents the data without dropout events. Cells were colored by CCPE-inferred cell cycle stages. **(b)** The impact of dropout rate on the performance of CCPE, Cyclum and CYCLOPS evaluated by clustering metrics, RI, ARI, NMI, ACC using simulated datasets. The methods are marked with different colors.

### Detection of G1 arrest in Nutlin-treated cancer cell lines using CCPE

To further assess the performance of CCPE, we applied CCPE to the cancer cell datasets with or without treatment with nutlin. Nutlin is a MDM2-p53 inhibitor [35] and can induce cell cycle arrest [36]. One dataset was from the cells treated with vehicle DMSO and another one is from the same cells treated with nutlin [37]. The cells used in culture were a cancer cell mixture with seven TP53 WT cell lines and seventeen TP53 mut cell lines. As shown in **Figure 6a**, TP53 WT cells were in red circle and cells were colored by CCPE-infered cell cycle stages. Compared with the cells in the control group treated with DMSO, CCPE successfully detected an increase in the number of TP53 WT cells in the G1 phase treated by Nutlin. (**Figure 6a**). We screened out the data of the seven TP53 WT cell lines and calculated the cell number ratio in each cell cycle phase. We found a significant increase of G1-phase cells, which confirmed that Nutlin can elicit a pronounced G1 arrest in TP53 WT cells compared with the untreated control (**Figure 6b)**. We also applied Deseq2 to identify the DEGs associated with CCPE-inferred cell cycle stages. It is obvious that some of the top ten enriched cell cycle pathways of these DEGs are associated with cell cycle, such as regulation of cell cycle progression and cell cycle G2/M checkpoint (**Figure 6c)**. The enrichment analysis of DEGs further illustrates the accuracy of CCPE in estimating cell cycle stages and the reliability of CCPE to successfully detect G1 arrest in nutlin-treated TP53 WT cells.

**Figure 6.**
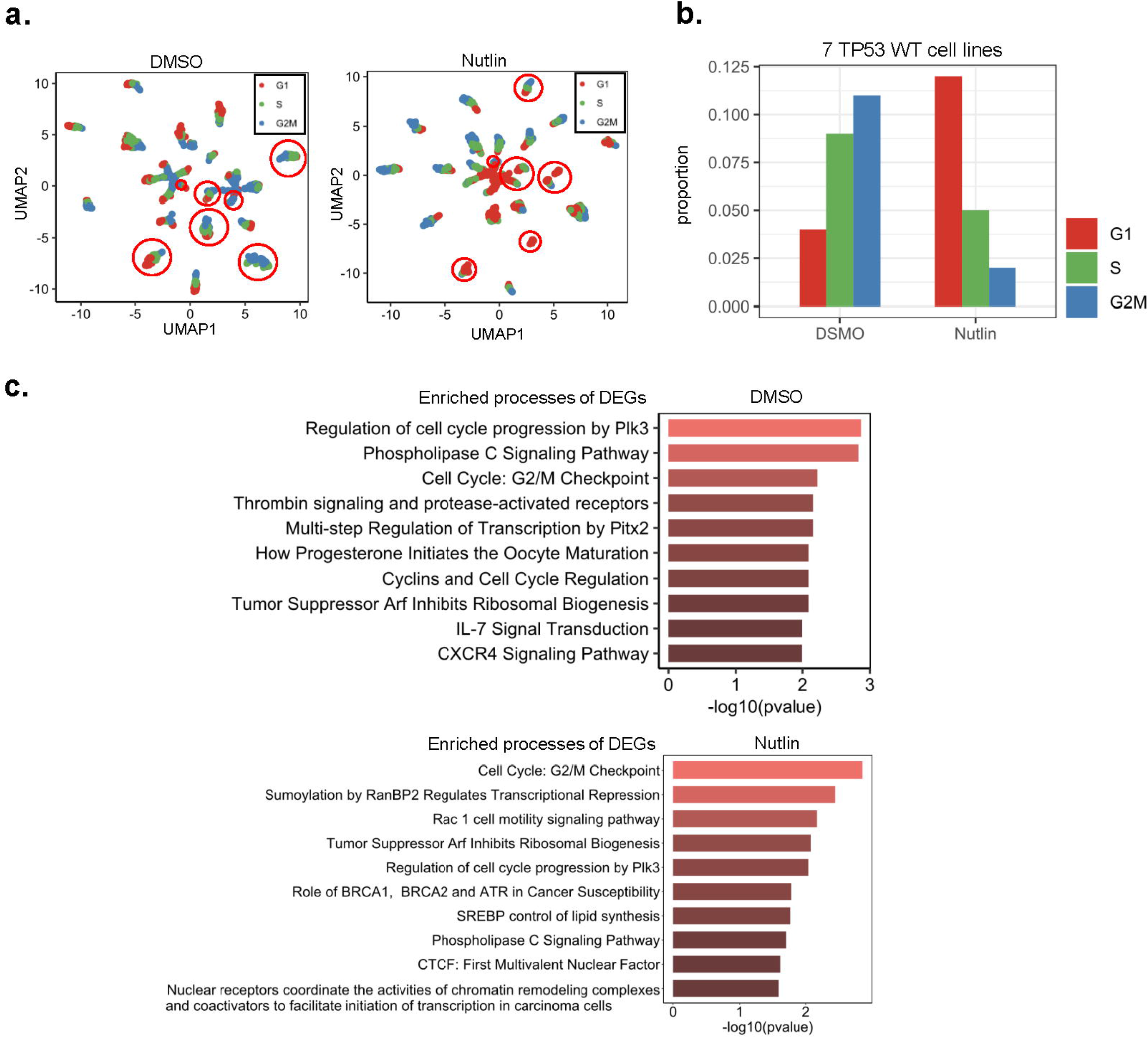
Validation of CCPE prediction accuracy using scRNA-seq datasets from cells treated with cell cycle disruptors. **(a)** UMAP plots for 24 cancer cell lines treated with DMSO and nutlin, separately. Cells were colored by CCPE-inferred cell cycle stages and cells in red circle are TP53 WT cells. **(b)** CCPE-inferred cell cycle stage distribution in nutlin-treated and control groups. **(c)** Top ten biological processes of of DEGs identified from CCPE-inferred cell cycle stages.

### Cell cycle effect removal from scRNA-seq data

The cell cycle is a major source of bias that introduces large within-cell-type heterogeneity, which obscure the differences in expression between cell types [38]. Investigating effective removal of cell cycle effects is important in such a situation. We use the murine multipotent myeloid progenitor cell line 416B dataset [39] to access the utility of CCPE in removing cell cycle effect. We compute the percentage of variance explained by the CCPE-inferred cell cycle stages in the expression profile for each gene. Genes with high percentages are regarded as cell cycle-related genes and are removed from the dataset [40]. We found that there is a small effect caused by cell cycle in the 416B dataset. Cells with two phenotypes can be distinguished but those with induced CBFB-MYH11 oncogene expression were separated from the raw data. After removing cell cycle effect using four methods, CCPE, Cyclum, Seurat and ccRemover, CCPE can separate two phenotypes correctly and the viariation between two phenotypes were more pronounced compared with raw data. Cyclum and Seurat can divide cells into two groups, but do not correspond to the expected phenotypes. CcRemover performs the worst, not being able to distinguish between the two phenotypes at all **(Figure 7)**.

**Figure 7.**
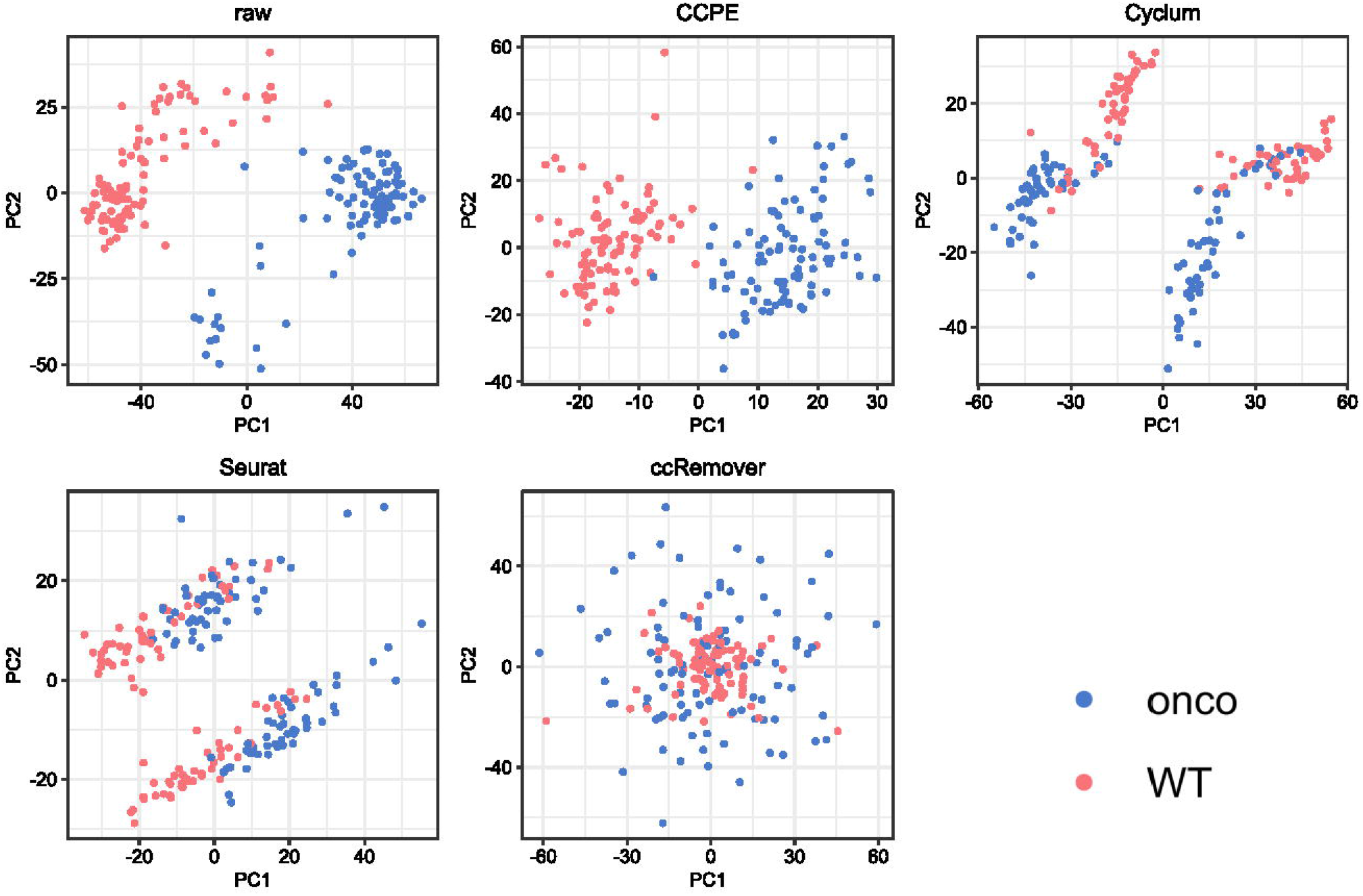
PCA plots of the 416B dataset, generated before and after removing cell cycle effect using CCPE, Cyclum, Seurat and ccRemover. Each point corresponds to a cell that is colored by oncogene induction status.

## Discussion

Pseudotime analyses of single-cell RNA-seq data have been increasingly used to determine the latent pattern of dynamic processes experienced by cells, such as the cell cycle [41]. We defined cell cycle pseudotime to describe the progression through the entire cell cycle process. Clustering is a common step to group cells into different cell cycle stages, learning gene expression pattern within different subgroups [42]. It is widely used in single-cell transcriptomics workflows. However, cell cycle is a dynamic process that gene expression varies between cells not subgroups. Cell cycle pseudotime analysis attempts to characterize such differences by project cells along a continuous process rather than dividing cells into discrete clusters [21]. In this study, we developed a novel cell cycle pseudotime estimation method named CCPE to accurately characterize cell cycle timing for single cell RNA-seq data. CCPE learns a discriminative helix with two dimensions to characterize circular process in cell cycle and one dimension to symbolize the pseudotime of cells along the cell cycle process. This is a kind of task in manifold learning, a strategy to learn the intrinsic structure of complex and high-dimensional data. We used alternating structure optimization to fit the best helix from scRNA-seq data. The parameters were optimized in the iterative transformation of high and low dimensional spaces. Discriminative information of cells in the same cell cycle phase were taken into consideration during the optimization process. Although CCPE is designed to predict cell cycle pseudotime, it can convert the pseudotime into discrete cell cycle stages through a Gaussian mixture model. Our strategies have enhanced the prediction accuracy of CCPE in predicting the cell cycle pseudotime and discreting cell cycle stages compared with currently existing methods.

Gene selection is recommended during data pre-processing of CCPE. Since single cell RNA sequencing data suffers from many sources of technical noise [43]. Some cell cycle estimation methods only use cell cycle genes, such as cyclone, Seurat and reCAT. Cyclone applied thousands of cell cycle gene pairs to determine the cell cycle phases of cells [6]. While in Seurat, only small number of S phase marker genes (43) and G2M phase marker genes (54) are used to identify the cell cycle [20]. The semi-supervised algorithm, reCAT, used 378 cell cycle genes listed in Cyclebase3 [44] to get the expression matrix, while other genes were excluded based on the risk of adding noise to the model [22]. From their performance on real scRNA-seq datasets, it is difficult to exactly figure out how many cell cycle genes are sufficient to accurately predict the cell cycle. On the other hand, there are some genes which are influenced by cell heterogeneity and partially contribute to the cell cycle. If these genes were completely ignored, then additional noise would be introduced to the cell cycle prediction. Therefore, we recommended to use a sophisticated approach called “dpFeature” to select differentially expressed genes during pre-processing of CCPE. “dpFeature” discovers the important ordering genes from the data, rather than relying on cell cycle marker genes from literature.

We assessed the performance of CCPE in estimating cell cycle pseudotime and various applications using both simulated and real scRNA-seq datasets. Even though CCPE is an unsupervised algorithm, we compared it with both knowledge-based and other unsupervised algorithms, including cyclone, Seurat, Cyclum, CYCLOPS and reCAT. Peco is not included in the comparison since fluorescence imaging is required with scRNA-seq to measure cell cycle phase. The mESCs Quartz-Seq dataset is widely used in various cell cycle studies [6, 22, 45]. We compared the perform CCPE with several algorithms in characterizing the cell cycle pseudotime using mESCs Quartz-Seq dataset. CCPE not only captured the right order of three cell cycle phases, but also separated them very well as expected. Additionally, correlation analysis shows the genes highly correlated with CCPE-inferred cell cycle pseudotime are G2/M phase marker genes. Gaussian mixture model in CCPE was applied to estimate discrete cell cycle states. We calculated ten clustering metrics on real datasets and our results indicated that CCPE had an outstanding performance compared with Cyclum, CYCLOPS and reCAT. We also tested the performance stability of CCPE in predicting cell cycle stages when the number of cells and genes in the dataset is small. Enrichment analysis showed that the DEGs identified by CCPE-inferred cell cycle stages had more connection to the biological processes related to cell cycle pathways. To exam the performance of CCPE in analyzing the datasets with high dropout events, we generated three simulated datasets with different dropout rates. CCPE had a strong capability to predict cell cycle states on the data with 25.6% dropouts. When the dropout rate increase, the performance of CCPE was reduced, but still outperformed than Cyclum and CYCLOPS. Cyclone, Seurat and reCAT require preliminary gene list and cannot be applied to simulated datasets, so we did not compare with these methods on our generated datasets. To further validate the performance of CCPE, we used CCPE to analyze the datasets collected from mixed cells treated with a cell cycle perturbation reagent nutlin. Nutlin, a selective MDM2 inhibitor and MDM2 is a negative regulator of the tumor-suppressor gene TP53. McFarland *et al.* [37] used Seurat to identify the cell cycle phase of each cell and concluded that Nutlin elicits rapid apoptosis and cell cycle arrest in G1 phase exclusively in the TP53 wild-type cells compared with the untreated cells. CCPE successfully caught the G1 arrest induced by nutlin in TP53 WT cells. Differential gene expression analysis further validated the accuracy of CCPE in estimating cell cycle phases. Removing cell cycle-related genes inferred by CCPE enhances differences between two phenotypes for 416B dataset.

CCPE is based on the manifold projected on 3D helix, which contains both circular process and linear axis. In future studies, we plan to use CCPE in the study of mechanisms involving both linear and nonlinear components, such as cell heterogeneity by the combination of cell cycle modeled by nonlinear component and cell types modeled by linear component from scRNA-seq data. In addition, CCPE uses soft clustering method rather than hard clustering assignment to account for cell cycle discriminative information, making smooth transitions between cell states between different cell cycle phases. The soft clustering algorithm favors clusters with cells from multiple datasets and preserves discrete and continuous topologies while avoiding local minima that might result from maximizing representation too quickly across multiple datasets [46]. The application of soft clustering in CCPE inspires the potential ability of CCPE to predict the cell cycle of datasets with different experimental and biological factors, which is what we plan to investigate next.

## Materials and Methods

### Datasets

We used both simulated datasets and real datasets to evaluate the performance of CCPE.

#### Simulated scRNA-seq datasets

We simulated three datasets with different dropout rates (25.6%, 51.1% and 68.8%) using the simulation model in CIDR [34]. Each simulated dataset contains three cell stages, representing G1, S, G2/M phases. One hundred fifty cells and 20180 genes were generated for each simulated dataset by setting parameters *N*=3, *k*=50 in *scSimulator* function of CIDR package. Different dropout rates (25.6%, 51.1% and 68.8%) are achieved by setting the dropout level parameter *ν* equal to 6.5, 9, and 12, respectively. Higher *ν* means a higher level of dropouts.

#### mESCs Quartz-Seq data

The mouse embryo stem cells (mESCs) were sequenced by Quartz-Seq technology, a reproducible and sensitive single-cell RNA seq method [18]. We used this dataset to analyze the accuracy of CCPE-predicted pseudotime. The mESCs Quartz-Seq data has 35 mouse embryo stem cells, including 20 cells in G1 phase, 7 cells in S phase and 8 cells in G2/M phase. mESCs Quartz-Seq data is available from Gene Expression Omnibus (GEO) with GEO Series ID GSE42268.

#### H1 hESCs scRNA-seq data

To compare the performance of CCPE and Cyclum on the data with different gene and cell sizes, especially the data with small number of genes and cells. We randomly subsampled the scRNA-seq data from human embryo stem cells (GSE64016). This dataset consists of 247 cells and 19084 genes. We selected 7 gene sizes, ranging from 50 to 600 genes, and 5 cell sizes, ranging from 10 to 100 cells. Each data with a specific size was sampled 10 times for fair evaluation. Normalized expected counts were provided in this dataset and the cell cycle phases of 247 cells were identified using Fluorescent Ubiquitination-based Cell Cycle Indicator (FUCCI).

#### E-MTAB-2805 mESCs data

This scRNA-seq dataset of mouse embryo stem cells were generated by Buettner *et al.* [45]. The dataset was downloaded from https://www.ebi.ac.uk/arrayexpress/experiments/E-MTAB-2805/. The cells were stained with Hoechst and sorted using FACS for respective cell-cycle fractions (G1, S and G2M phase). Two hundred eighty-eight mouse embryo stem cells were sequenced using HighSeq 2000 sequencing system. After cell selection, 279 cells were used in CCPE to analyze differentially expressed genes.

#### Mix-seq dataset

This dataset consists of two 10x single-cell RNA-seq data from nutlin-treated cells and control group. A mixed culture of 24 cell lines were treated with either dimethyl sulfoxide (DMSO) or nutlin. This dataset was downloaded from https://figshare.com/s/139f64b495dea9d88c70. Nutlin is known to elicit cell cycle arrest exclusively in cells expressing wild-type (WT) TP53s [37]. Thus, seven cell lines expressing WT TP53 were used in CCPE to character the cell cycle effect of a cell cycle perturbation.

#### 416B cell line

The dataset contains two 96-well plates of 416B cells (an immortalized mouse myeloid progenitor cell line) [39], processed using the Smart-seq2 protocol [47]. The 416B cell line includes wild type phenotype and induced CBFB-MYH11 (CM) oncogene expression. This dataset was downloaded from https://www.ebi.ac.uk/arrayexpress/experiments/E-MTAB-5522/.

### Normalization and Pre-processing

For the mESCs Quartz-Seq data and E-MTAB-2805 data with FPKM and TPM expression levels, and other datasets with read counts for expression levels, we normalized the single cell RNA-seq datasets by taking log 2 transformation with a pseudo count 1 as

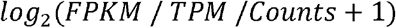

Gene and cell selections are recommended for the data pre-processing in CCPE. ‘dpFeature’ is a powerful and general approach, for unsupervised feature selection [48]. We used dpFeature to select differentially expressed genes between cell cycle phases. Low-quality cells enriched with low expressed genes were not included in CCPE.

### CCPE model

#### Learning a Helix in Reduced Dimension

We suppose scRNA-seq data X with N cells and D genes lies in the high dimension. In CCPE, we considered the linear projection *z* = *f* (*X*) = *W^T^X* to infer the embedded expression profiles Z_dxN_ (d<<D) from X and the reversed linear projection is *X* = *f*^−1^(*Z*) = *WZ*, where *W* ϵ *R*^*D*×*d*^ and *W^T^W* = 1. We construct a circular helix 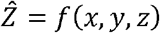 in 3D dimension to get the best fit of Z, a circular helix of radius *a* and slope *ν*/*a* (or pitch 2*πν*) is described as follows for cell *i*

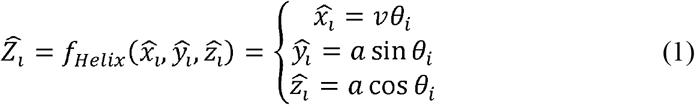

Then we formulate the following object function to obtain the reduced dimension via learning a helix

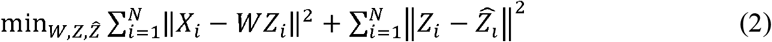

Where *X* = [*X*_1_, *X*_2_, … , *X_N_*] ϵ *R*^*D*×*N*^ is the scRNA-seq data, *W* = [*W*_1_, *W*_2_, … ,*W_d_*] ϵ *R*^*D*×*d*^ is an orthogonal set of *d* linear basis vectors *W_l_* ϵ *R^D^*, *Z* = [*Z*_1_, *Z*_2_, … ,*Z_N_*] ϵ *R*^*d*×*N*^ is represented by the embedded expression profiles of *X* in low-dimension 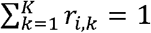 contains fitted points of *Z* on a circular helix with the same dimension as *Z* and 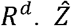 is the coordinates of cell *i* projected on the circular helix.

Furthermore, cells in the same cell cycle phase should cluster together on the helix, so we consider the clustering objective into the optimization problem as below

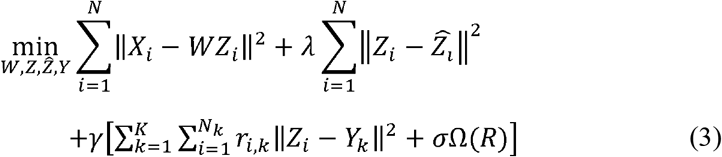

where *N_k_* is the number of cells in cluster *k*. *r_i,k_* is the weight of soft clustering based on the assumption of *K* clusters. Ω(*R*) is the negative entropy regularization and *σ* > 0 is the regularization parameter for Ω(*R*). *λ* > 0, *γ* > 0 are parameters that indicate the importance of each component of the objective function. The solution of *r_i,k_* in terms of 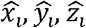 and formula of Ω(*R*) are described in [49] as the following

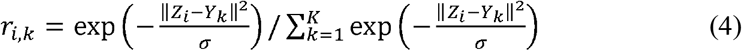

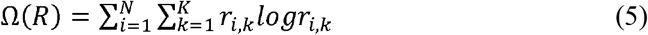

### Optimization of CCPE

We optimize the object function (3) using alternating structure optimization, which has been successfully applied to several optimization problems [50]. We divide the parameters to be optimized into two parts {*W,Z,Y*}and 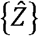 and solve one group by fixing the other group alternatively until convergence.

Firstly, we optimize {*W,Z,Y*} by fixing 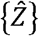. Given a known helix, we can see 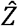 as a constant matrix *C* ϵ *R*^*d*×*N*^. After simple matrix manipulation, function (3) with respect to {*W,Z,Y*} can be rewritten as the following optimization problem

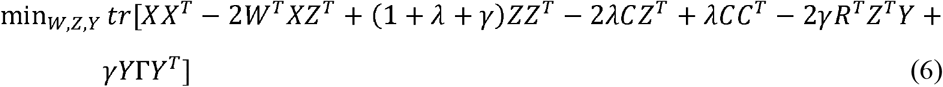

Where Γ = *diag*(1^*T*^*R*) and *R* is the weight matrix of soft clustering. Set *L* equals formula (6) and the first derivative of *L* with respect to *Y* to zero

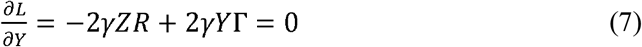

Then we get the optimization of *Y* as

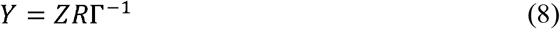

Substituting *Y* into *L* and set the first derivative of *L* with respect to *Z* to zero, we can get the optimization of *Z* as

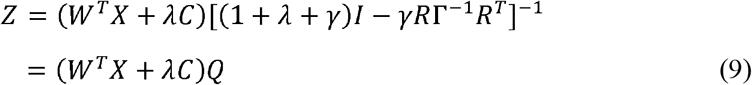

where *Q* = [(1+ λ + *γ*)*I* − *γRΓ*^−1^*R^T^*]^−1^ and the inverse of [(1+ λ + *γ*)*I* − *γRΓ*^−1^*R^T^*] exists. Similarly, substituting *Z* into *L* the objective function becomes the following optimization problem

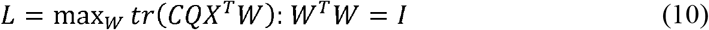

This is the constrained quadratic problem which has the closed-form solution [51] of *W* as follows

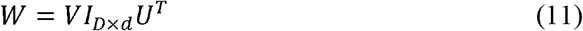

where *V* is an *D* × *D* unitary matrix, *U* is an *d* × *d* unitary matrix, and *UΣV^T^* is the singular value decomposition of matrix *CQX^T^*, *Σ* is an *d* × *D* rectangular diagonal matrix with non-negative real numbers on the diagonal.

Secondly, given *W*, *Z* and *Y* we can obtain 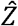 easily by solving the following curve fitting problem

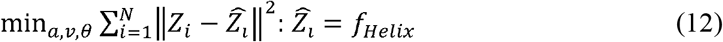

Overall, the optimization process of the problem (3) is given in **Algorithm 1**.

**Algorithm 1.**
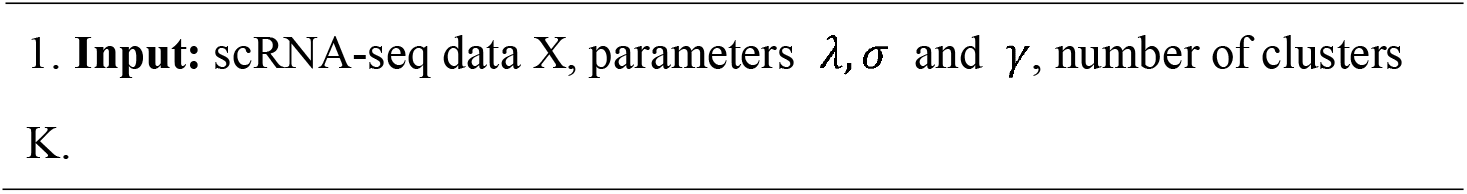
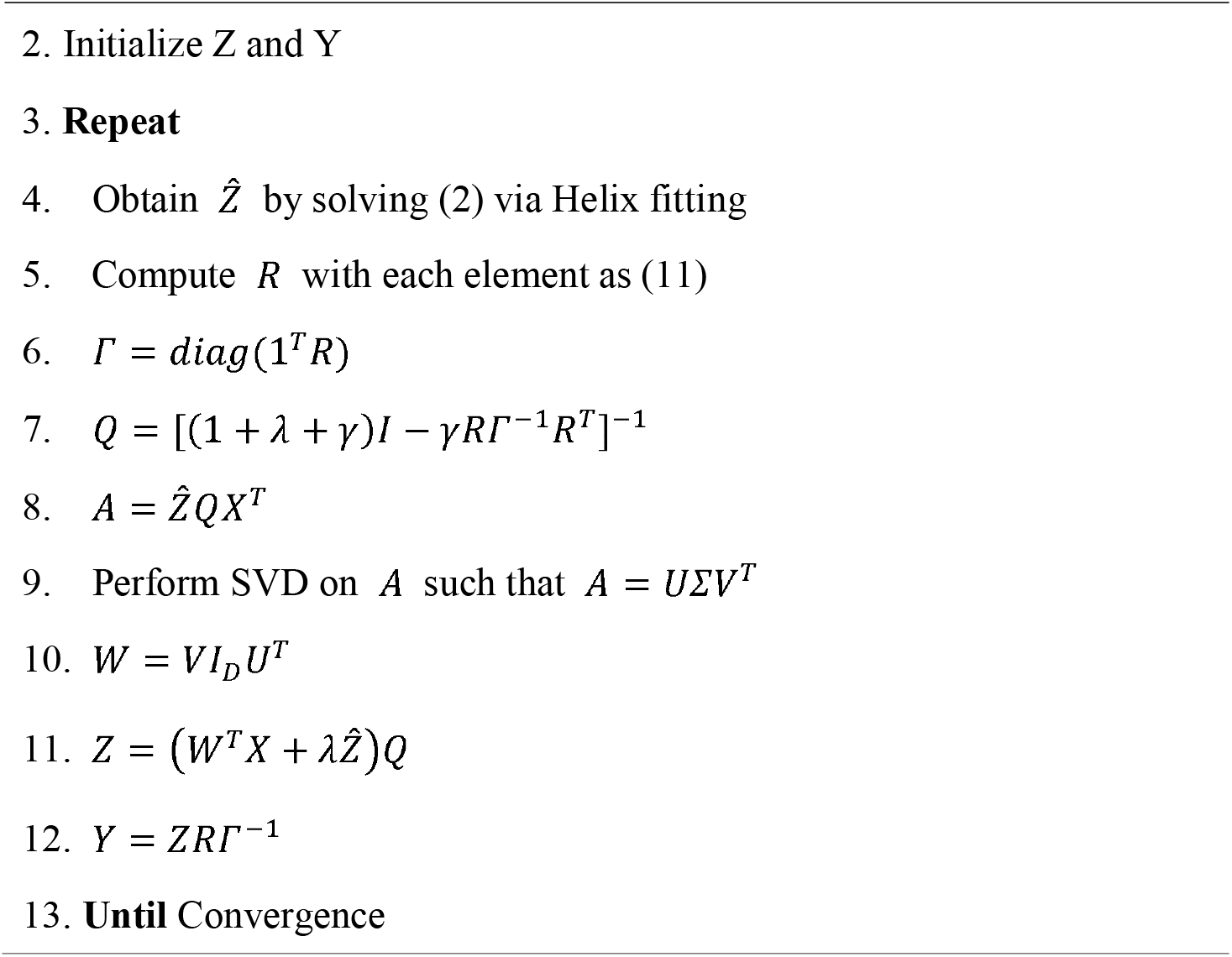

## Availability of Data and Software

CCPE is an available tool in the GitHub repository (https://github.com/LiuJJ0327/CCPE). The source of the datasets we used is described in **Materials and Methods**.

## Conflict of Interest

The authors declare that the research was conducted in the absence of any commercial or financial relationships that could be construed as a potential conflict of interest.

## Funding

This study was supported by the National Institutes of Health [R01GM123037, U01AR069395-01A1, R01CA241930 to X.Z]; The funders had no role in study design, data collection and analysis, decision to publish or preparation of the manuscript. Funding for open access charge: Dr & Mrs Carl V. Vartian Chair Professorship Funds to Dr. Zhou from the University of Texas Health Science Center at Houston.

## Acknowledgments

We thank the members of the Center for Computational Systems Medicine (CCSM) for valuable discussion and suggestions.

## Notes

### Competing Interest Statement

The authors have declared no competing interest.

